# Trypanosomes Modulation of Rotational Motility from Swimming to Network-Threading Propulsion in Confined Environments

**DOI:** 10.1101/2025.11.03.686241

**Authors:** Narges Jamshidi Khameneh, Timothy Krüger, Patryk Nienaltowski, Yves Emery, Dmitry A. Fedosov, Marco Polin, Markus Engstler

## Abstract

*Trypanosoma brucei* (*T. brucei*) is a protozoan parasite that lives extracellularly in the body fluids of its hosts. In mammals, these environments include the vascular system and the interstitial spaces of various organs, such as the skin and adipose tissue. How the parasite disseminates within the host remains largely unresolved. It is clear however, that parasite motility plays a central role. The unicellular, eukaryotic flagellate is a versatile microswimmer, like bacteria or sperm cells, albeit structurally far more complex than these classical model systems. In addition to possessing a flagellum attached alongside a strongly polarised cell body, the parasite is capable of swimming both forwards and backwards. The trypanosome must be capable of navigating effectively even under extreme physical and mechanical constraints. It can do so in the mammalian host with only one main morphotype. This means that the cell is mechanically adapted to motion in diverse challenging microenvironments. To address how, we study the parasites in different viscoelastic regimes up to conditions mimicking tissue confinement. Next to quantitative high speed video microscopy, we employ digital holography microscopy, yielding three-dimensional subcellular resolution. We detail the mechanical reaction of the flexible cell body with its uniquely attached flagellum to increasing confinement and show how the trypanosomes are able to maintain their motile capabilities to spread in dense tissue.

## Introduction

African trypanosomes are unicellular parasites that cause severe diseases in humans and livestock. The salivarian species causing important neglected tropical diseases like sleeping sickness, are transmitted by the tsetse fly (1). Although they have been studied for over a century, only in recent years has some attention shifted to the physics of their locomotion (2– 6).

One impediment is the morphological complexity, compared to the intensively studied free flagellar motility of sperm cells or other selected microswimmers (7). The flagellum is extensively attached to the cell body. Only a relatively short portion extends beyond the anterior pole of the cell. This extremely unusual structural configuration raises questions on the evolutionary advantage of such a morphology.

Another difficulty is that the parasites rarely reveal their true swimming capacities under cell culture conditions. The bloodstream forms (BSF) of *Trypanosoma spp*. thrive extracellularly in the circulation and tissues of infected vertebrates (4,8), where conditions deviate from those in standard cell culture. Forwards or backwards travelling waves along the single flagellum move the cell through fluids. The near simultaneous wave propagation in both directions results in the tumbling, non-progressive movement of the cell typically observed in low viscosity culture media. In contrast, under physiological flow prevailing in the bloodstream *T. brucei* is stimulated to more unidirectional and thus progressive swimming in the name-giving, corkscrew-like rotational fashion (6). The basic motility mechanism is used by all trypomastigote cell types analysed so far, in the tsetse fly (9) and by BSF of various African trypanosome species (10).

In mammals, the environments trypanosomes must navigate include the vascular system and the interstitial spaces of various organs. Different tissue-specific forms have been described (11–21), but the motile behaviour was not explored in any detail. While the parasites display considerable morphological diversity in the tsetse fly, the forms found in the mammalian host are relatively uniform in shape. This suggests a structural capacity to flexibly adjust to changing surroundings purely by mechanical means. In the bloodstream, trypanosomes are exposed to constantly shifting flow velocities, shear forces, and extreme cellular crowding. In addition, in narrow capillaries, the cells are highly constrained, as the vessel diameter is only marginally larger than that of the parasite itself. In lymphatic vessels, different flow conditions prevail, while lymph nodes present additional physical barriers. The interstitial space of tissues and organs is dominated by the extracellular matrix (ECM), which is composed of mesh-like structures of collagen and elastin, as well as highly viscous polymers, including proteoglycans. All of these extracellular compartments are physically interconnected and in constant exchange (4,22). Thus, the parasite faces a continuum of hydrodynamic and elastic resistances.

In summary, the extracellular fluid space of mammals is a system of mechanically extremely hostile environments. Thus, it is unsurprising that African trypanosomes are the only infectious agents that populate all of them. The question of how the parasite disseminates within the host remains largely unresolved, although it is clear that parasite motility plays a central role. The trypanosome must be capable of navigating effectively even under extreme mechanical constraints, making it a rich model organism for studying the physically complex interactions of a eukaryotic cell with a large range of challenging microenvironments.

To address this, we focused on how trypanosomes adapt cell morphology and swimming performance to conditions of increasing viscosity and confinement mimicking what is expected to occur in tissues. Such analyses have been an area of intense research, but have mainly been conducted with much simpler cells, such as bacteria and sperm. Whereas the theoretical foundation of micro-swimming in low viscosity and Newtonian fluids is extensive and solid (7,23,24), the impact of complex fluids and spatial confinement on the motility of microswimmers has only recently opened a rich field of interest in biological fluid mechanics and active matter, with potentially important implication on the biology of swimming microorganisms (25–27).

We detailed trypanosome swimming behaviour using two- and three-dimensional analyses with subcellular resolution. We observed distinct changes in flagellar waveform and subtle yet effective adjustments in the fusiform morphology of the parasite. We observed the trypanosomes′ ability to maintain net swimming speed across different viscosities, up to conditions of confining surroundings that force the cells to reverse movements and explore different routes. This provides, for the first time, insight into the refined, purely mechanical adaptations that trypanosomes employ to cope with rapidly changing microenvironments.

## Results and Discussion

To assess the influence of environmental viscosity on persistent forward swimming of *T. brucei* bloodstream form cells (*BSF*), we analysed cell motility under both low (1 mPa*s, 5 mPa*s) and high (250 mPa*s, 1000 mPa*s) dynamic viscosity (μ), using polymeric methylcellulose solutions. The high-viscosity conditions were used to mimic the increasing physical confinement that trypanosomes encounter in dense extracellular matrices, prevalent in body tissues. The aim was to provide a simplified yet tuneable model system to investigate if and how *T. brucei* modulates its motility in response to increasing environmental resistance and structural complexity. At high viscosities the polymer solutions are in the dense entangled regime (> 40 mPa*s for methylcellulose (28)), which means that typical gaps between polymer chains resemble well the mesh size of gel-like networks. In denser polymer solutions, cells will become increasingly confined and viscoelastic effects will play a larger role.

As an reference, μ = 5 mPa*s approximates the viscosity of blood and has previously been shown to increase the persistence of BSF swimming (6). The viscosity µ = 250 mPa*s is in the range of high viscosity media used for motility analysis of sperm, which simulate midcycle cervical mucus (29). The high viscosity of 1000 mPa*s is comparable to pure glycerol at room temperature, and in this range viscoelastic effects can have varying effects on microswimmers (26). This viscosity is higher than that of media usually used for sperm motility analysis. It should be noted that body fluids with viscosities of around 1000 mPa*s can be found in joint cavities or in certain mucus layers. Neither microenvironment is populated by trypanosomes.

We analysed two cell culture-adapted *T. brucei* lines with differing viscosity requirements for growth: *MITat 1*.*6*, maintained in standard medium (μ = 1 mPa*s), and *AnTat 1*.*1*, maintained in high viscosity medium (μ = 250 mPa*s). The reason for these requirements is unknown, so we asked if the motility behaviour of the two cell lines changes differentially when subjected to environments of varying viscosity. We show here the results for the pleomorphic strain AnTat 1.1, which is a wildtype strain capable of progressing through the complete developmental cycle in flies and mammals. The comparison to the typical culture strain MITat 1.6 shows very similar results (Fig. S1). We conclude that forward motility is not a factor in the AnTat cell′s viscosity requirement. It has been proposed before that the necessity for high-viscosity medium is probably due to effects during cell division (30).

### Flagellar Wave Parameters Adapt to Viscosity

To investigate how flagellar dynamics adapt to different viscosities, we recorded videos of persistently forward swimming cells with 100 frames per second (fps), providing a temporal resolution of 10 ms. This allowed us to quantify the flagellar movements during propulsion at the level of individual flagellar beats (∼18-25 Hz) (31). From each time-lapse series we selected images corresponding to the initial bend of each oscillatory wave, as well as those in which the planar wave assumed a horizontal position within the focal plane (Fig. 1b-e, t = 0 ms). Using these images, the amplitude and wavelength of the anterior wave were measured. Using the sequences of images (Fig. 1b-e, t = 0 - x ms), the beating frequency and straight-line velocity were determined. By analysing the image sequences beat for beat until the cell had completed a full rotation around the axis of progressive movement, we also quantified the rotational frequency and the translocation per rotation, defined as the swimming distance during a single 360° rotation, (Fig. 1b-e, t = x ms).

**Fig. 1.**
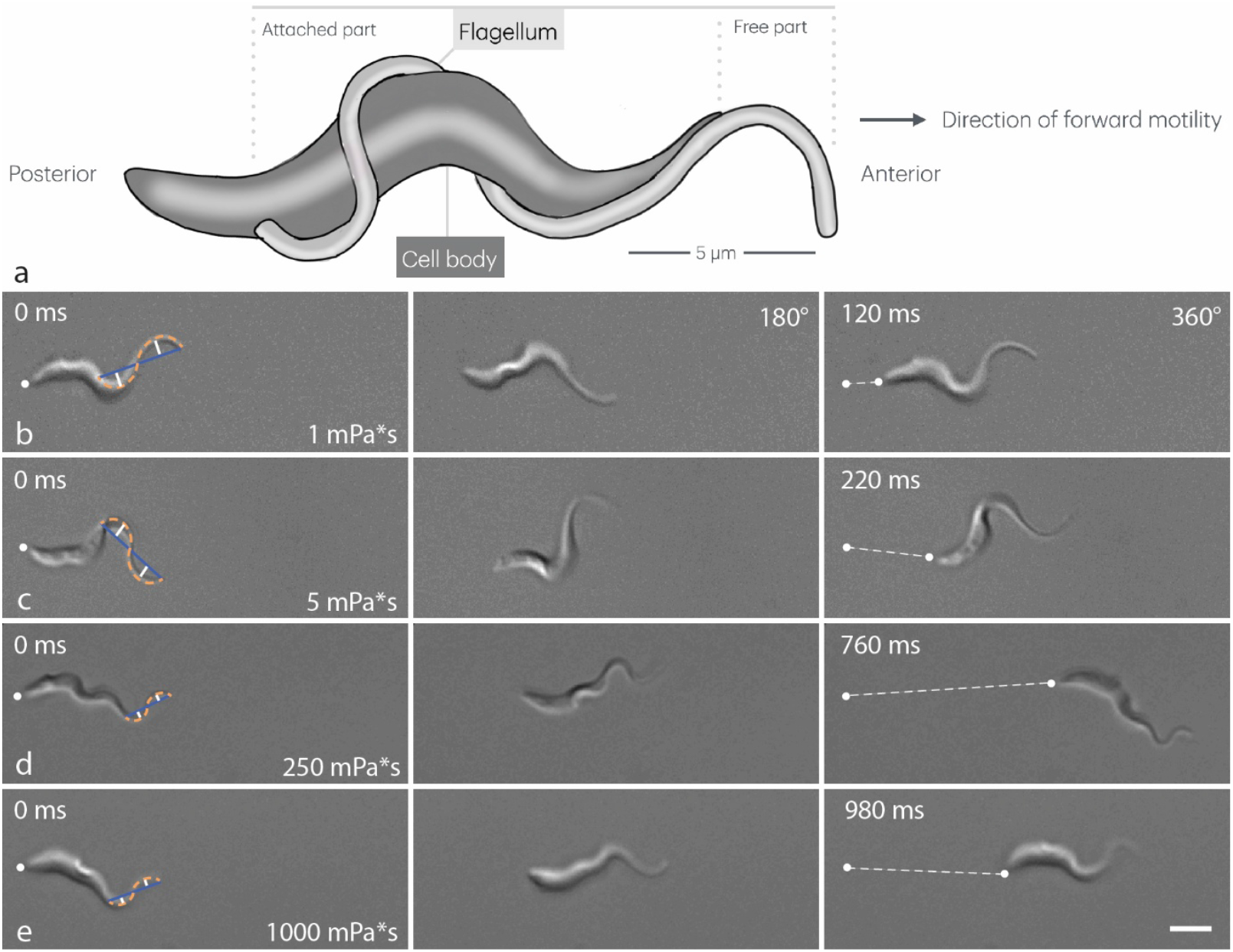
*T. brucei* persistent forward swimming during one rotation in different viscosities. **a)** Schematic of trypanosome morphology showing attachment of flagellum to the cell body and forward swimming direction. **b)** 1 mPa*s **c)** 5 mPa*s, **d)** 250 mPa*s, **e)** 1000 mPa*s. Each row shows three snapshots of the same field of view from videos recorded at 100 fps (Videos S1-S4). **Left**: Timepoint selected as t_0_ at the beginning of a flagellar beat. The image is chosen to show the flagellar wave oscillating in the image plane. The anterior-most wavelength is marked (wavelength: blue line, bending flagellum: dashed orange curve, amplitudes: white lines). The posterior tip is used to measure swimming distances and speed (white dot). **Middle**: Images of each cell rotated 180° while swimming to the right. Note the inverse curvature of the posterior (left) end of the cell body and the anterior-most (right) wavelength of the flagellum. **Right:** Cells after a full 360° rotation around its longitudinal axis (note the same body configuration as in b). The swimming speed was measured as the straight-line velocity between the localisations of the posterior tip (white dots, the dashed line represents the translocation per rotation or “lead”). Scale bar: 5 µm.

Our analysis revealed that all three wave gait parameters, amplitude (A), frequency (f) and wavelength (ʎ), were significantly reduced at higher viscosities (250 and 1000 mPa*s) compared to lower viscosities (1 and 5 mPa*s) (Fig. 2a-c). At viscosities of 250 mPa*s and above, the mean values of these parameters were approximately halved relative to their values in low viscosity (A (5 mPa*s) = 1,5 µm, A (250 mPa*s) = 0,8 µm; f (5mPa*s) = 22 Hz, f (250 mPa*s) = 12.9 Hz; ʎ (5 mPa*s) = 8.9 µm, ʎ (250 mPa*s) = 5.5 µm). Note that the flagellum has a fixed length and a reduction in wavelength leads to an increase in the number of waves, from approximately 1,5 at low viscosities to 3 - 4 at higher viscosities. This change is clearly visible when comparing Figs. 1b and c to Figs. 1d and e. However, precise quantification along the posterior part of the cell is difficult due to the chiral attachment of the flagellum and the attenuation of the wave amplitude in this region.

**Fig. 2.**
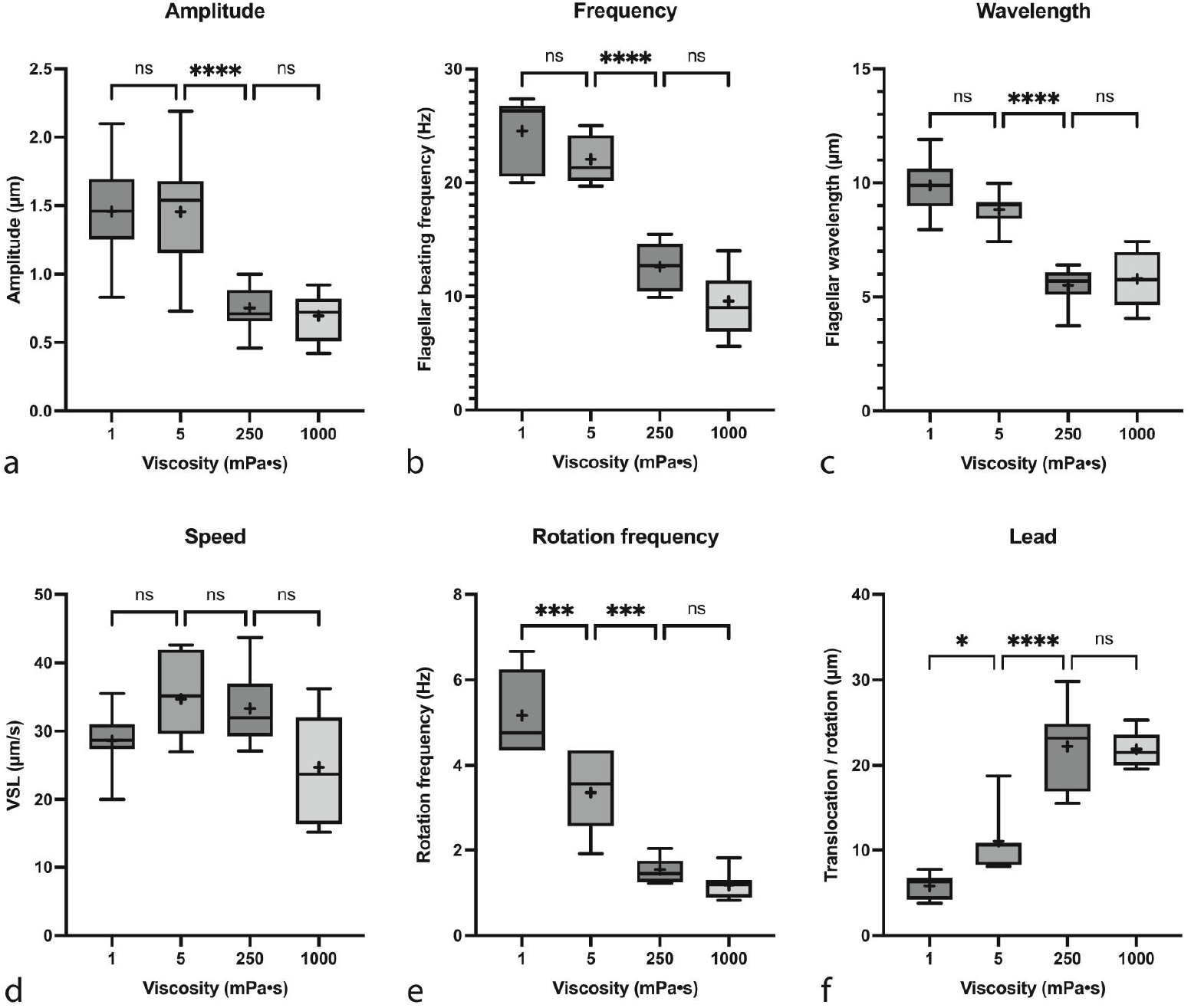
Quantification of *T. brucei* AnTat 1.1 forward swimming parameters. **a)** Amplitude of the anterior-most wavelength as measured during beating in the image plane (white lines in Fig. 1). **b)** Flagellar beating frequency measured as number of anterior-initiated waves during time periods of uninterrupted swimming. **c)** Wavelength of the anterior flagellum (blue lines in Fig.1). **d)** Swimming speed, measured as straight line velocity of the posterior tip (dashed white lines in Fig.1). **e)** Rotational frequency, measured as the number of uninterrupted 360° rotations around the longitudinal axis per time unit. **f)** Lead, measured as the translocation per 360° rotation (dashed white lines in Fig. 1). Box plots show the mean (+) and median (line) values within the 25th to 75th percentile range as boxes and the min to max values as whiskers (n=7 cells and 2-4 rotations for each parameter). Significance was analysed by ordinary one-way ANOVA (****: p < 0,0001, ***: p<0.001, **: p<0.01, *: p<0.05, ns: p>0.05).

These wave parameters are critical for the establishment of realistic trypanosome models (2,3) and for modelling propulsion of slender filaments, as described by resistive force and slender-body theories, which have been successfully applied to the flagellar propulsion in sperm cells and other microswimmers in Newtonian fluids (7,24,32–34).

Despite the pronounced changes in flagellar waveform across different viscosity conditions, the swimming velocity of the trypanosomes remained relatively stable. The average swimming velocity was 28.6 µm/s in 1 mPa*s and 22.6 µm/s in 1000 mPa*s, with the highest velocity observed in 5 mPa*s (34.7 µm/s) (Fig. 2d). Note, that across the entire spectrum, these speeds stay above the theoretical speed of 20 µm/s required for efficient antibody removal from the cell surface (35). These results indicate that the 2D wave parameters alone cannot fully account for the preservation of *T. brucei* swimming velocity at high viscosity.

Although the rotational frequency decreased in higher viscosities (Fig. 2e), indicating higher drag forces acting on each movement of the cell, the effective translocation per rotation increased significantly (Fig. 2f). This suggests that the rotational swimming mechanism enables the trypanosome to adapt to confined environments by adopting a flatter helical swimming trajectory, thereby enhancing directional propulsion along the longitudinal axis (Fig. 3c, d). Regarding the eponymous auger-like motion of the trypanosome and comparing it with the rotational progression of a screw, this effect corresponds to an increase in the screw’s lead, that is, the axial distance travelled during one revolution.

**Fig. 3.**
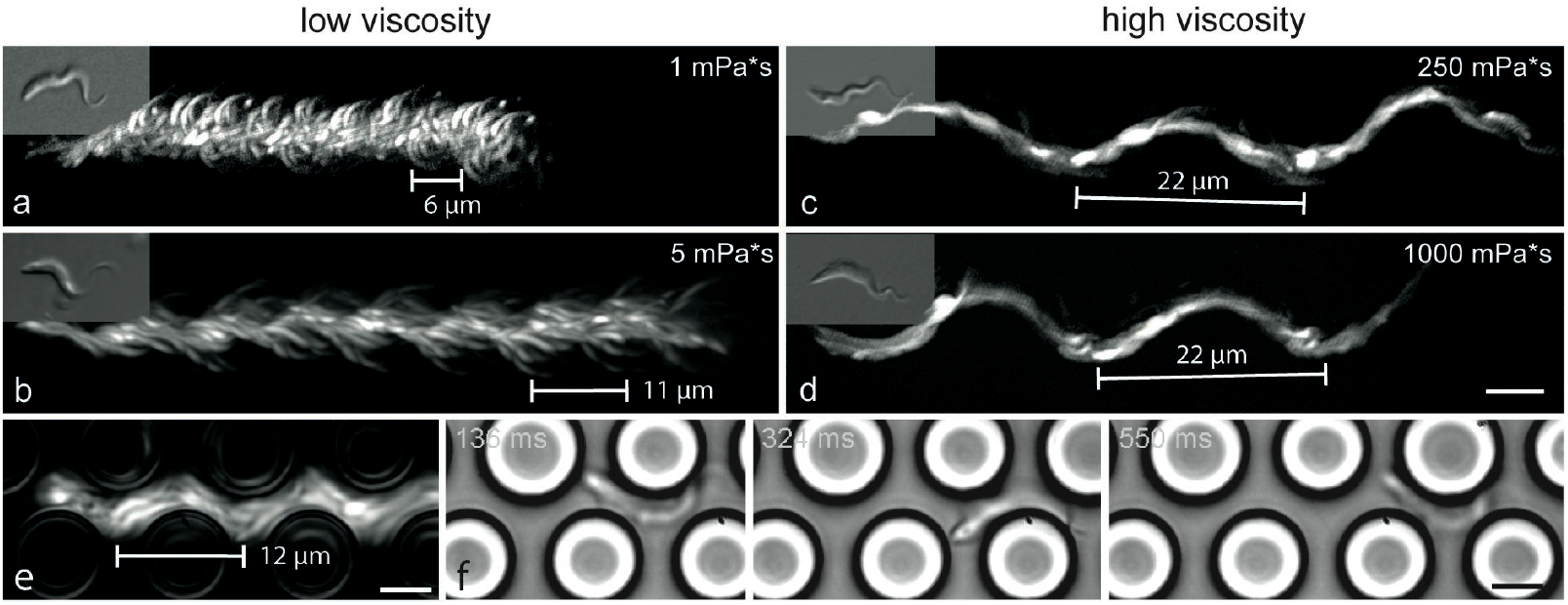
Adaptation of trypanosome trajectories to viscosity and solid resistance. **a)** Rotations with high frequencies and low translocation result in broad and straight trajectories in low viscosity. **b)** In 5 mPa*s the trajectory appears stretched as the lead almost doubles and swimming speed increases. **c)** In high viscosity surroundings the trajectories appear smoothed, flattened and wavy as the cells rotate smoothly around the longitudinal axis with a high lead. Velocity is constant despite higher resistance. **d)** In even higher viscosities the body does not stretch further, keeping the lead at maximum value and slowing slightly due to drag. **e)** Trypanosome swimming at 1 mPa*s viscosity in a PDMS pillar array (pillar diameter: 8 µm, spacing: 4 µm, data modified from (6)). **f)** Still images of the cell projected in e from Video S6, depicting angles 0°, 180° and 360° of one rotation. Images a-d are maximum projections of the cell shown in the corresponding inset. The time spans of the projected sequences in a-d are the same, as to depict the relative speeds of the cells. Measured distances represent the translocation per rotation. Projections of a-d correspond to supplemental video sequences in Video S5. Scale bars: 5 µm (Inset magnification = *0,5).

In contrast to the wave parameters, the rotational parameters showed significant differences between 1 and 5 mPa*s (Fig. 2e, f). As these low viscosity values are well within the Newtonian range, this indicates that here, only viscosity affects rotational behaviour and there is no additional influence of elasticity. As rotational swimming behaviour is directly correlated with the directional progression of microswimmers (36), this suggests that effective viscosity similarly influences the trypanosome locomotion, regardless of whether it arises from large physical obstacles (e.g. red blood cells), or from the presence of polymers with defined hydrodynamic radii (e.g. methylcellulose). This equivalence has previously been shown by the increased swimming persistence of trypanosomes under both conditions (6). Notably, similar effects, albeit based on different mechanisms, have been shown in other microswimmers, providing a surprisingly simple explanation for the increase in their swimming speed in complex fluids (25,27,37).

A direct comparison with the trajectory of a trypanosome swimming in a PDMS-pillar array at 1 mPa*s, showed that the lead is increased to about the same length as the swimmer in 5 mPa*s (Fig. 3e). The cell is mechanically restricted to the space between the pillars and the translocation per rotation (12 µm) is adapted exactly to the diameter of the obstacles (8 µm) plus the spacing between the pillars (4 µm), the parameters corresponding to the situation in the bloodstream (6). Note, that to test the equivalent situation for higher viscosities, the pillar spacing would need to be reduced further to reduce amplitudes to the submicrometric range (Fig. 2a). This is technically very difficult to realise, but would represent a smooth transition to continuous solid boundaries anyway, environmental constrictions as in the simulations in (3).

We did not observe further significant changes in the wave parameters when viscosity was increased from 250 to 1000 mPa*s (Fig. 2a-c, e, f). However, the mean speed decreased from 32,7 µm/s to 22,6 µm/s, which might indicate that the trypanosomes have reached the limit of their morphological adaptation for forward propulsion.

### Digital - holography reveals changes in cell bending under different conditions

The above results highlight the critical role of 3D rotational dynamics in maintaining effective and persistent motility under mechanical constraints. The observed changes in wave gait are closely interdependent with the elastic properties of the trypanosome body to which the flagellum is attached, and vice versa. To directly analyse the three-dimensional changes during rotation, we recorded persistently swimming cells under identical conditions using digital holographic microscopy (DHM). For an analysis of body bending, we tracked the central axis of the cell over a full rotation for swimmers in both low viscosity (1 mPa*s) and high viscosity (250 mPa*s) environments (Fig. 4). Additionally, the positions of the flagellar tip and posterior end were tracked to characterise swimming trajectories and rotational behaviour (Fig. 5). This approach allowed us to resolve the three-dimensional dynamic adjustments in body shape and motion patterns in response to environmental changes.

**Fig. 4:**
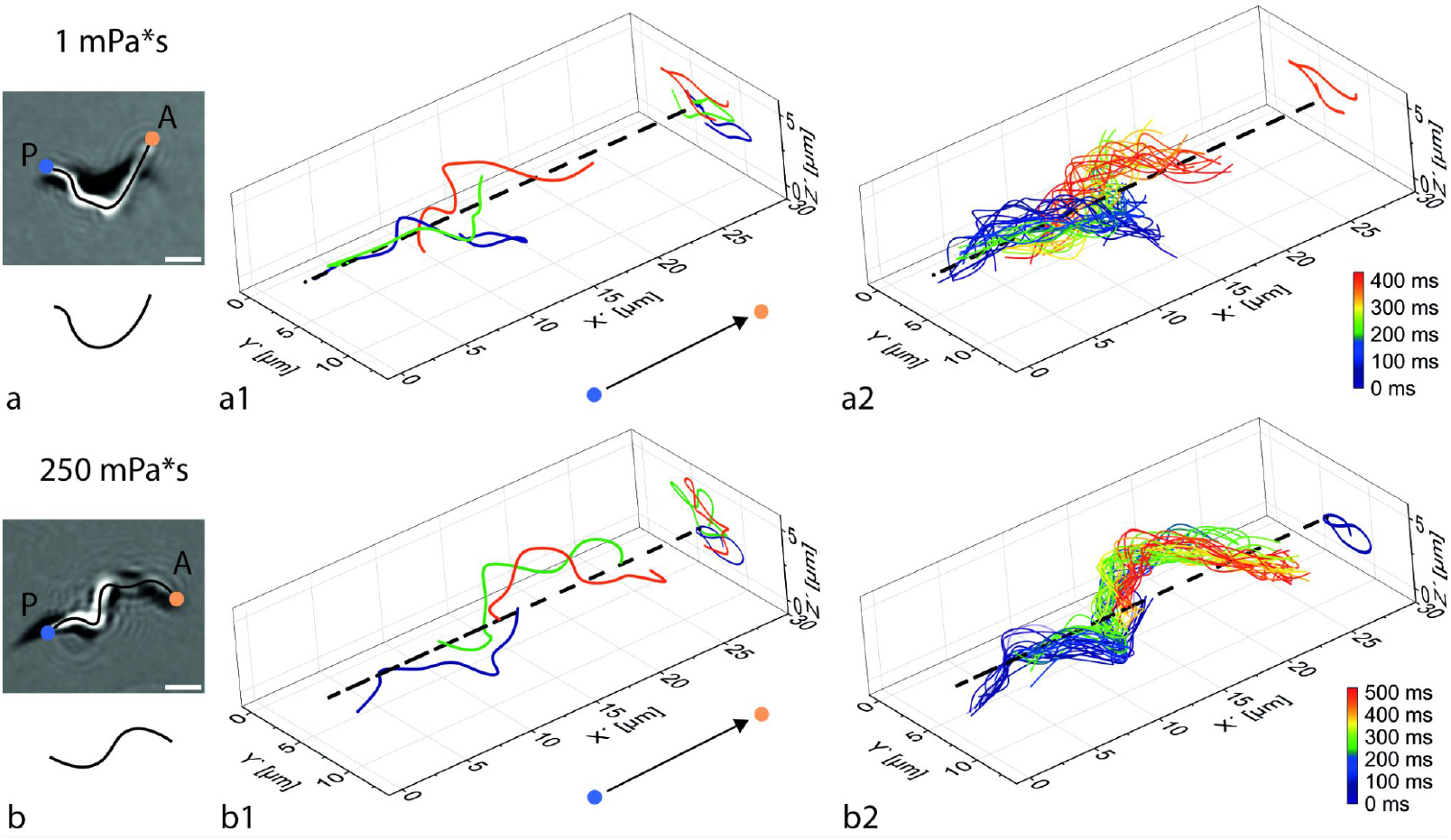
3D **c**ell body bending dynamics during one rotation in low and high viscosities. **a-a2)** Low viscosity (1mPa*s). **b-b2)** High viscosity (250mPa*s) **a, b)** Projected images of original DHM data used to trace the cell body axes (black lines). Anterior and posterior tips are marked with an orange and a blue dot, respectively. Scale bar: 5 µm. Note: The line below the image is a smoothed schematic to illustrate the principal change in bending; compare to *yz*-projections in a2, b2. **a1, b1)** Selected 3D traces of the cell bodies of one trypanosome during one rotation each. The *yz*-projections in a1 show the half helix turn the body traces typically constitute around the axis of movement (open c-shape), whereas the projections in b1 show the full helical turn represented by a closed s-like shape. The *x*-axis is oriented parallel to the axis of rotation (dashed black line, equivalent to progressive swimming axis), as determined by principal component analysis (PCA). The swimming direction is indicated by the arrows. **a2, b2)** Complete set of time colour-coded body traces in 10 ms steps, visualising the 3D trajectory of the bending cell. Only one of the selected traces each is shown in *yz*-projection as plotted in a1 and b1 for better visibility. Rotatable graphs are linked as html files for a2 and b2 (Supplemental material).

**Fig. 5:**
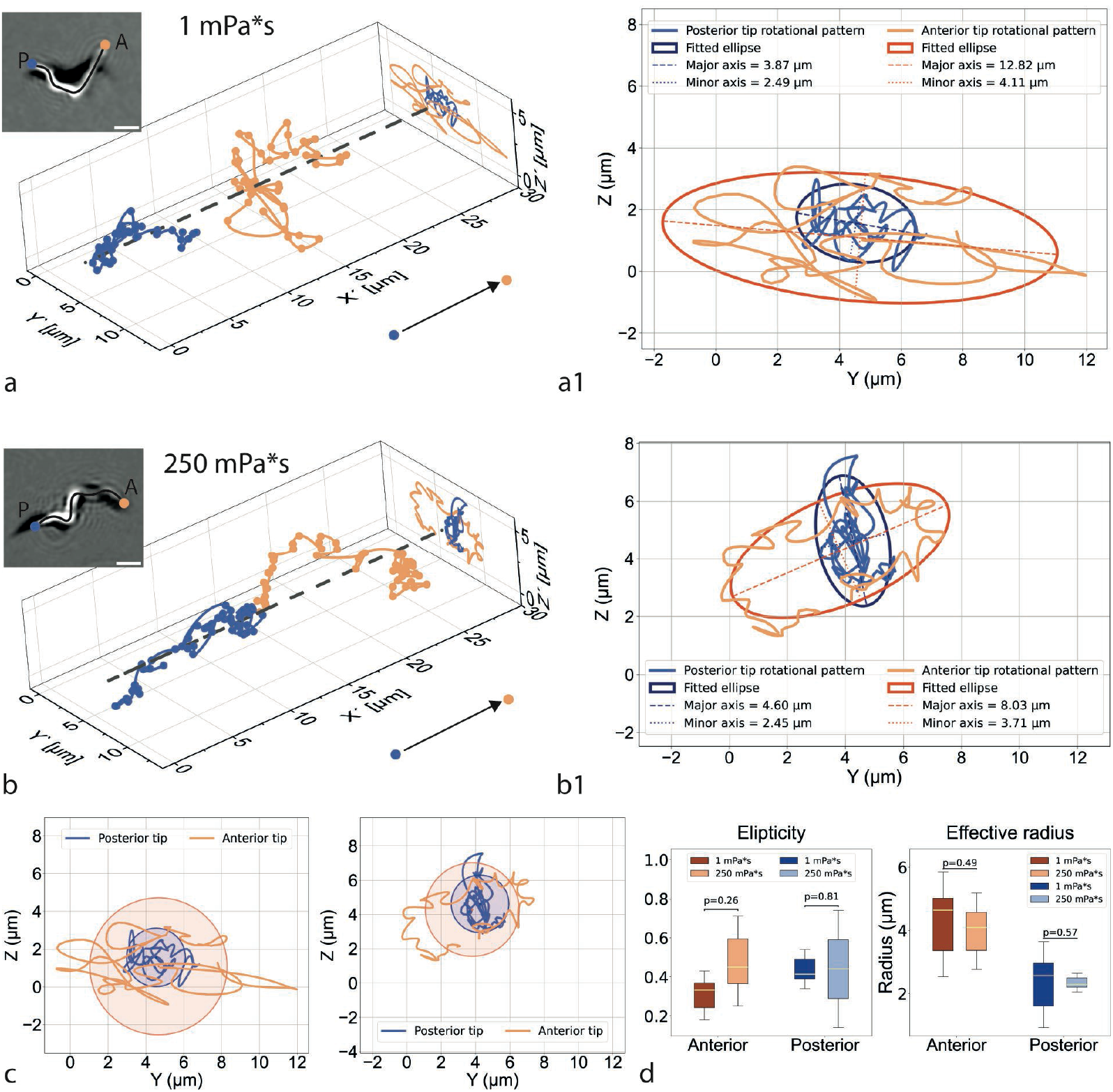
Rotational behaviour quantified by 3D single point tracing. **a, b)** The movement of the exemplary cells used in Fig. 4 (insets, scale bar: 5 µm) were tracked by tracing the posterior tip of the cell body (blue dots) and the anterior tip of the flagellum (orange dots) in low viscosity (a, 1mPa*s) and high viscosity (b, 250 mPa*s). The axis of rotation was defined by the first principal component of PCA using all data points (Coordinate system as in Fig. 4). The points were connected using splines to visualise the progressive and rotational movement along and around the axis of rotation. The tracks were projected on the *yz*-plane. Rotatable graphs are linked as html files for a and b (Supplemental material). **a1, b1)** Magnified view of the *yz*-projections in a, b showing the gyrating behaviour of the tips during rotation. The points were fitted to ellipses which represent the flattened helical trajectories of the cells. **c)** The data points were used to fit circles to calculate the effective radii of the helical trajectories. **d)** The ellipticity calculated from the ellipse axes from several rotations showed no significant difference between low and high viscosities. The effective radii show the clear difference between the large exploratory radii of the more flexible anterior tips and the smaller radii of the posterior tips. The radii show no significant difference between low and high viscosities.

The body traces show the helical bending of the cell around the progressive swimming axis. In low-viscosity conditions the swimmer typically exhibits only a half turn of a helix at a given position, making the projections of the traced body line appear as roughly C-shaped lines rotating around the swimming axis (Fig. 4a). In contrast, under high-viscosity conditions, the cell body constantly forms a complete helical turn, resulting in a rather S-shaped configuration (Fig. 4b). This full helical twist produces a prolonged, flattened cell shape, which likely contributes to the more effective translocation per rotation in high-viscosity medium, akin to a screw that maintains traction with the surrounding medium, while advancing more smoothly. In contrast, the half pitch configuration observed in low-viscosity swimmers, results in greater bending away from the progressive axis, which reduces propulsion efficiency, but the lower viscous drag enables the parasites to compensate through higher amplitudes, beating and rotation frequencies. (Figs. 2, 3, Video S5).

The tracing data provided a three-dimensional perspective on the characteristic trajectories (Fig. 4a2, b2) previously seen in two-dimensional projections (Fig. 3), completing the missing information on the symmetry of the bending and twisting dynamic cell morphology. To quantify the rotational behaviour, we analysed the positions of the posterior cell pole and anterior flagellar tip using single point tracking (blue and orange dots in Fig. 5a, b, respectively). These points were tracked over one rotation and plotted to reconstruct the trajectories around the swimming axis (Fig. 5).

As expected, the plotted tip trajectories (Fig. 5a, b) follow the helical trajectories visualised by the body-axis traces (Fig. 4a2, b2). The projection of both traces results in roughly elliptical envelopes of different size (Fig. 5 a, b). This is a representation of the 3D beating as an elliptical helicoidal waveform, which is typical of experimentally and theoretically described 3D waveforms in sperm cells, and which arises due to asymmetries in the actuation and morphology of the cells (38,39). We fitted ellipses to these projected traces for quantification of the 3D beating envelopes (Fig. 5 a1, b1). The anterior flagellar tip traces the maximal extent of the envelope around the cell trajectory, as the most widely oscillating point of the cell, whereas the posterior tip is restricted to a consistently smaller volume. This is projected in the smaller radii of the ellipses (blue in Fig. 5).

Note, that the tips show gyrational movement around the main helical trajectory of the cell. Due to the flexibility of the trypanosomes’ body, both ends are free to exhibit lateral movements in all directions, deviating from the main helical trajectory (the anterior end more so than the posterior). These wobbling movements are the result of the bending we described with the body traces above and are stronger in low viscosity medium. This gives rise to a more pronounced looping of the tip trajectory projections (Fig. 5 a-b1). These swimming patterns arise because of the asymmetry of the cell body and its attached flagellum, similar to the patterns seen in sperm cells, when flagellar beating is asymmetric relative to the sperm head. Although the resolution is not sufficient to characterise the trajectory pattern perfectly, it clearly resembles trace classifications like the spinning star trajectories of sperm cells (39).

The ellipticity of the waveforms (Fig. 5d) shows a fundamental constriction of movement in one direction of the cells′ moving coordinate system. This presumably has a largely viscosity-independent, structural cause. The chiral attachment of the flagellum to the cell body is maintained tightly through the flagellar attachment zone and most likely dictates a plane where freedom of oscillatory movement is lower, thus causing the trajectory to deviate from a perfectly circular helix.

Interestingly, although high-viscosity swimmers follow a more compact helical trajectory, the area of exploration around the swimming axis does not differ significantly (Fig. 5 c, d). This further illustrates the trypanosome′s capacity to sustain directional movement under mechanical constraint. Across a wide viscosity range, the system of flagellar wave modulation by mechanical forces, which controls the deformation of the attached elastic cell body, enables trypanosomes to sustain the same radius of exploration. At the same time, the swimming mechanism is adjusted via the gait parameters for continuous progressive swimming capability.

The effects of body elasticity on trypanosome progressive and rotational motility have recently been studied by mesoscale hydrodynamic simulations (2). The results showed the influence of body stiffness and flagellar attachment on progressive speed and rotational frequency under constant viscosity. These results define an optimisation of cell morphology regarding these parameters under a specific mechanical condition of the microenvironment. It will be interesting to evaluate the characteristics of these parameters, which are not experimentally measurable to date, in future simulations in environments of varying viscosity.

### Boundary effects at high viscosity lead to non-rotational circular swimming

So far, we have described the motility mechanism of persistently forward swimming cells. The analysed cells were recorded in the centre of fluid volumes of 10 µm height, ensuring that the cells had no direct boundary contact. The cells also showed the same motility behaviour in chambers with a fluid height of 250 µm. In culture medium with a viscosity of 1000 mPa*s we observed a striking change of swimming behaviour in the vicinity of the confining glass surfaces. The cells ceased rotating altogether but did not experience any changes in wave gait otherwise. As a result, the cells continued to persistently swim in the forward direction but were limited to a circular swimming trajectory (Fig. 6a, Video S7). In the 250 µm chambers, this behaviour was only observed in cells swimming in around 3 µm proximity of the cover slip.

**Fig. 6:**
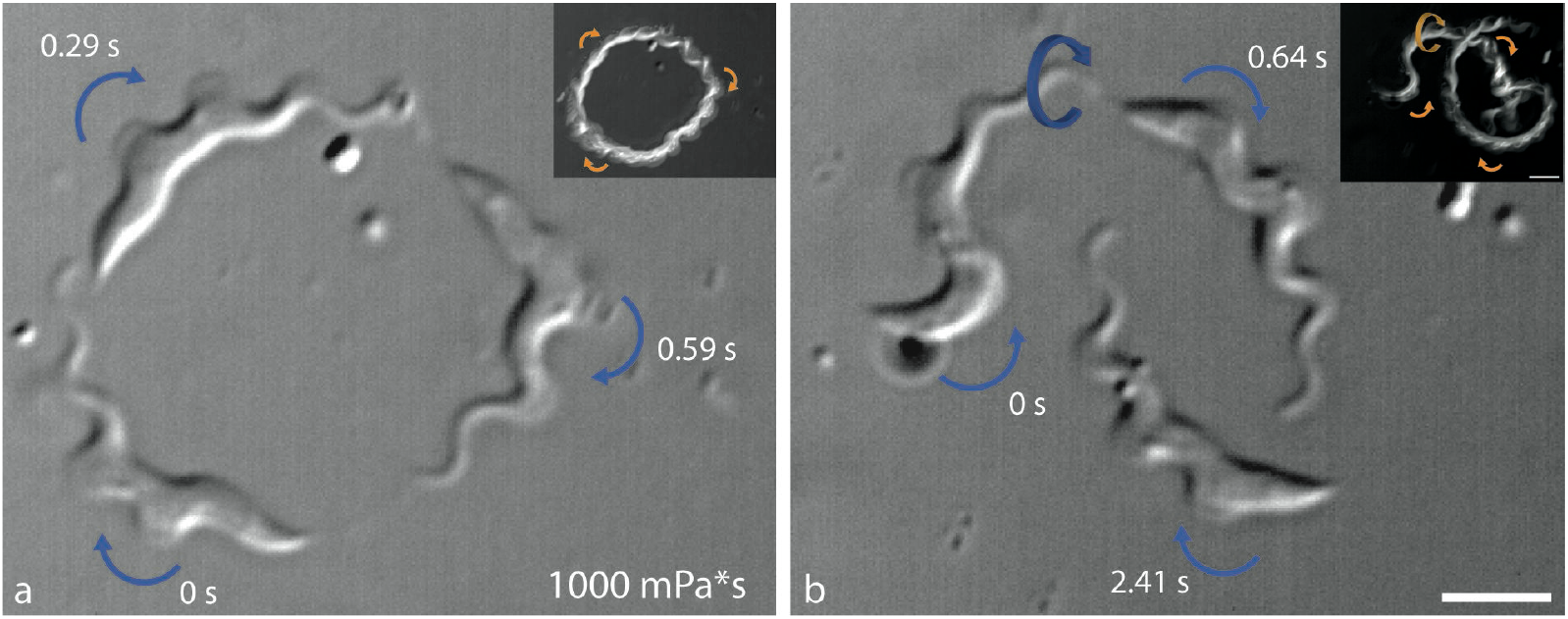
Circular, non-rotating swimming and switch to progressive swimming by rotation **a)** Persistent forward beating without rotation leads to circular swimming at 1000 mPa*s. **b)** Cell switching from a progressive helical trajectory to circular trajectories (indicated by 3D-arrow) as rotation is inhibited (Note, there is a short interrupting backward beat after t = 0.64s that shifts the circular trajectory to the right).Orange arrows indicate the swimming direction and the curvature of the posterior part of the cell at the depicted timepoints. Scale bars = 5 µm. Video sequences shown in Videos S7 and S8.

This clearly shows the necessity of rotational movement for progressive forward swimming in an asymmetric microswimmer, as has been well analysed for sperm cells (39–41). The switch from a 3D helical swimming path to a circular trajectory is also highly reminiscent of sperm cell behaviour close to glass surfaces (40,42). Further, the full ability of the trypanosome cell body to twist around its longitudinal axis is demonstrated. Together with the torsional changes shown in different viscosity regimes (Fig. 4), the entire ensemble of microtubule corset, flagellar attachment structures and flagellar axoneme can be seamlessly twisted from a curved, linear arrangement (Fig. 6b, Video S8) to the S-shaped helical structure seen in Fig. 4b.

The implications of the circular swimming for the *in vivo* behaviour of trypanosomes are unclear, but it is easily envisaged that the parasites could find themselves in a situation where they are trapped in very thin fluid layers of interstitial spaces between the highly viscous extracellular matrices of the tissue layers they are crossing. There is no swimming inhibition, and the cells can reverse to progressive movement as soon as they experience a change in the microenvironment, allowing them to rotate again. This behaviour was observed as a switch between progressive and circular swimming in the inhomogeneous methylcellulose (Fig. 6b, Video S8).

### Trypanosomes can navigate dense tissue-like networks by bending and beat reversal

We have proposed that the confinement imposed on trypanosomes by an increasingly dense network of polymers, causes the cells to adapt with a maximally stretched helical morphology that enables them to uphold a high persistent swimming velocity. We assume that this reaction is most likely due to elastic effects of the entangled polymer network confining the cells in the high viscosity solutions. We now wanted to know how the swimming behaviour changes with even higher viscoelasticity and confinement. Therefore, we recorded cells in methylcellulose solutions with a viscosity of 4000 mPa*s and in collagen gels.

In the 4000 mPa*s environment cells increasingly experienced interruptions of forward flagellar beats, making it virtually impossible to find uninterrupted persistent forward swimmers. Instead, more often a reversal of swimming direction was observed, as the trypanosomes apparently got stuck in the dense polymer network and reversed the flagellar wave progression from posterior to anterior, resulting in backward swimming (Fig. 7a, Video S9). By reversing the flagellar beat, the cells can change the direction of swimming and explore the surroundings in another direction (6). In low viscosity surroundings, the cells produce their trypanosome-characteristic tumbling behaviour in this way. In contrast, in the confining network a smoothly rotating and drilling motion was observed, where the parasite probes accessible space in the polymer mesh in three dimensions, frequently reversing along the same path. Wherever there was enough space to manoeuvre, the anterior flagellar tip bent to begin new forward beats in a different direction (Fig. 7a, Video S10).

**Fig. 7:**
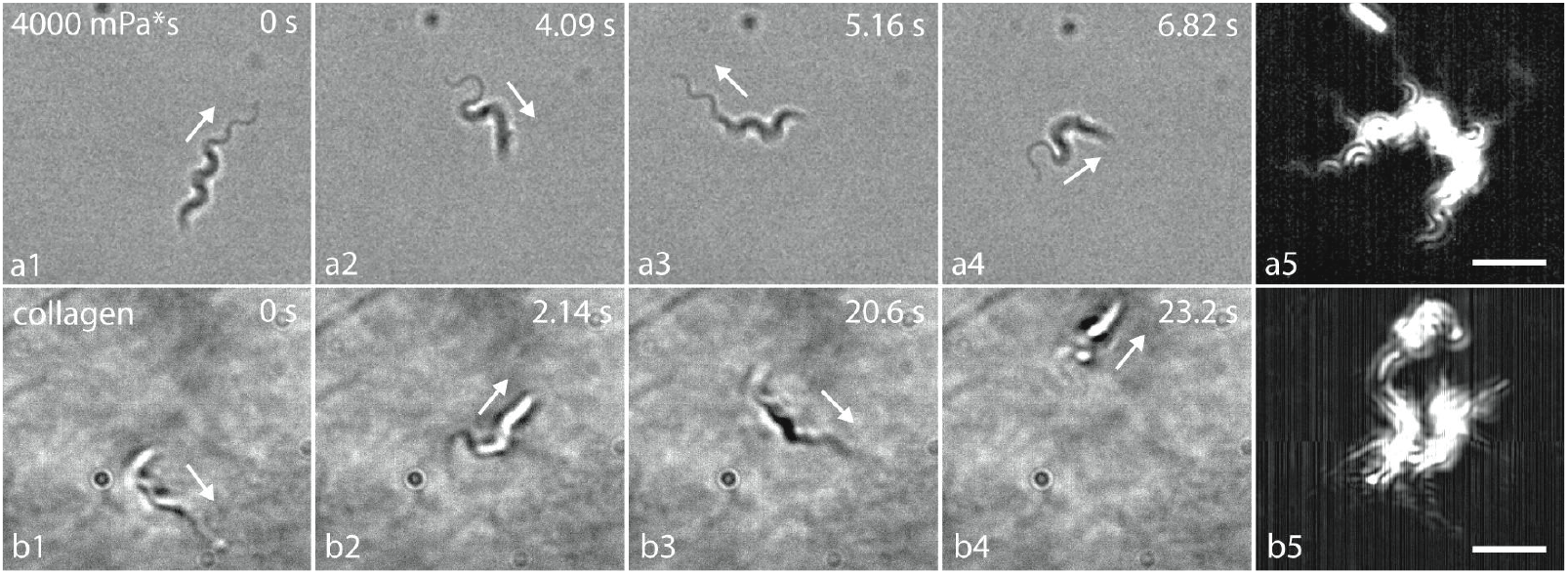
Trypanosomes traverse dense networks by reversals and flexible wriggling in three dimensions. **a)** Cell in 4000 mPa*s methylcellulose solution probing the field of view in several directions. Snapshots showing the extended waveform of high viscosity swimmers during forward movement (a1, a3) and a typical more curled-up waveform during backward movement (a2, a4). **b)** Equivalent probing of a volume of a collagen network (4%). Snapshots show the comparable type of movement as in a); The forward movement in the extended waveform (b1, b3) and the change to the more compact reversing shape (b2, b4). The dense meshwork mostly distorts the regular waveform seen in a) and reverse swimming is more prominent. The volume covered by the back-and -forth movements in a given period is highly variable and depends on the density of the immediate surrounding network. Scalebars 10 µm. Full video sequences are shown in Videos S9 and S10.

This was similar to the behaviour we observed after injecting trypanosomes into collagen gels (Fig. 7b, Video S10). In general, the cells were confined even more by the collagen network, making the waveforms appear disrupted and the cell shape crimped, as it writhed forwards and backwards through the tight and inhomogeneous fibrous meshwork. Whereas the majority of movements were forwards with reversals in methylcellulose, the collagen matrix caused the cells to move backwards more persistently between direction changes. In the dense network, the cells also showed more 3D-movement as the cell tested available space in all directions, showing the full extent of its flexibility.

Collagen is the main structural component of tissues and reconstituted extracellular matrices and thus the starting material to mimic biological tissue properties. The viscoelastic moduli are still lower than soft tissues (43), but we have observed the same type of probing motility during the dissemination of skin tissue form *T. brucei* in advanced artificial tissue models (16). The focus in that work was on proliferation of the tissue form populations, but the skin models will be instrumental for analysing the parasites′ dissemination. Thus, we believe that we can elucidate the swimming and the spreading mechanism of *T. brucei* in hosts, by bridging a range of confining microenvironments as proxies, from viscous fluids to viscoelastic soft tissues.

In summary, we describe the reaction of trypanosome dynamic morphology and the resulting swimming behaviour over a range of environments imposing increasing mechanical confinement on the cells. In Newtonian fluids, increasing resistance against flagellar deflections by increasing concentrations of polymers or the presence of immobile obstacles leads to an increase of swimming speed, more effective rotational movement and higher persistence. With increasing confinement, viscoelastic effects start to significantly negatively influence the wave gait characteristics essential for flagellar swimming. Nevertheless, the cells’ progressive swimming speed is not decreased to the same degree. Rather, the rotational motility is adjusted by a change of helicity of the flagellum and attached cell body, enabling the parasites to smoothly bore their way through thicker territory. We show the limit of this twisting adjustment and the conditions at which the cells are forced to stop rotating or get stuck abruptly, impeding progressive movement. Even in conditions of high confinement, the cells can find space to carry on swimming, by moving until rotation is possible again, or by switching to backward beating to reverse out of dead ends.

We believe the described adaptivity of the flexible trypanosome flagellum with the attached cell body to be a good description of the relevant motile behaviour in the vertebrate host system, especially for its indispensable penetration of tissues, e.g. skin for further transmission. Therefore, the described versatile mechanism is also of general interest for further research of mechano-receptive mechanisms in flagellated microswimmers.

## Material and Methods

### Cell culture

*T. brucei* BSF cell-lines *MITat1*.*6* and *MITat1*.*2* were cultivated in HMI-9 medium. Pleomorphic *AnTat1*.*1* were cultivated in 1.1% A4M methylcellulose (MC) in HMI-9 medium. All cells were grown at 37°C with 5% CO_2_. The cells were maintained in exponential growth phase at concentrations between 5*10^4^ and 5*10^5^ cells/ml.

### Viscous media

For motility analysis, media were prepared with varying concentrations of A4M methylcellulose (MC) in HMI-9 medium. Final concentrations of 0.4%, 1.1%, 1.5%, and 2% MC yield media with respective viscosities of 5 mPa*s, 250 mPa*s, 1000 mPa*s, and 4000*mPa&s. HMI-9 alone has a viscosity of 1 mPa*s.

Collagen type I from rat tails at a concentration of 8 mg/mL (kindly provided by Florian Groeber-Becker, University of Wuerzburg) was dissolved in 0.1% acetic acid and diluted 1:1 in HMI9 medium. The collagen solution was dispensed into tissue culture-treated 96-well plates and incubated overnight at 37°C.

For experiments, final cell concentrations of 8×10^5^ cells / ml medium were used. For MC solutions, cell motility was recorded in 5 μl medium between a microscope slide and a 25×40 mm coverslip. Alternatively, 80 µl of medium was pipetted into a Gene-Frame chamber (Thermo Fisher Scientific) that was then sealed with a coverslip to create a chamber of ∼250 µm height. For recording in collagen, 5 μl of medium were injected with a Nanofil syringe (34G needle, WPI) into each gel prepared in the well plates. All data was recorded at room temperature (23°C).

### Imaging

Widefield imaging was performed with an inverted microscope (DMI 6000B, Leica) equipped with a 40× (NA = 0.75) dry and a 63× (NA = 1.3) glycerol immersion objectives under differential interference contrast (DIC) illumination. Videos were recorded with an sCMOS camera (pco.edge, Excelitas PCO GmbH) at 100 fps.

### Digital holographic microscopy

3D data was recorded using a commercial off-axis digital holographic microscope (T-1000, Lyncée Tec). The system utilised partially coherent illumination at 666 nm, collimated for uniform sample illumination (44). Cells were recorded using 40× (NA = 0.75, HCX PL FLUOTAR, Leica) and 63× (NA = 1.3, HC PL FLUOTAR, Leica) objectives. Videos were captured at a frame rate of 100 fps. Holograms were reconstructed into *z*-stacks with an axial spacing of 0.4 μm using the Koala software (Lyncée Tec), supplemented with custom Python scripts for data analysis. The localization method was user-guided: for each frame, the operator identified the most in-focus plane, from which the corresponding *x, y, z* coordinates were extracted.

## Data analysis

2D video analyses were carried out using ImageJ. 3D visualisations and corresponding analyses were conducted using Python or OriginPro23 (OriginLab). Statistical analysis and plotting were performed with Prism 10 (GraphPad).

## Acknowledgements

Many thanks to Sandro Käser and Thorsten Ochsenreiter (University of Bern) for support during the holography experiments. The authors acknowledge support of the European Training Network PHYMOT (Horizon 2020 Marie Sklodowska-Curie grant agreement No 955910; N.J.K., P.N., M.P. and M.E.). M.P. acknowledges the fact that IMEDEA is an accredited ‘María de Maeztu Excellence Unit’ (grant CEX2021-001198, funded by MCIN/AEI/10.13039/501100011033). M.E. acknowledges support by the German Research Council (DFG) through the DFG priority programme 2332, ‘Physics of Parasitism’.

## Supplemental Material

**Fig. S1.**
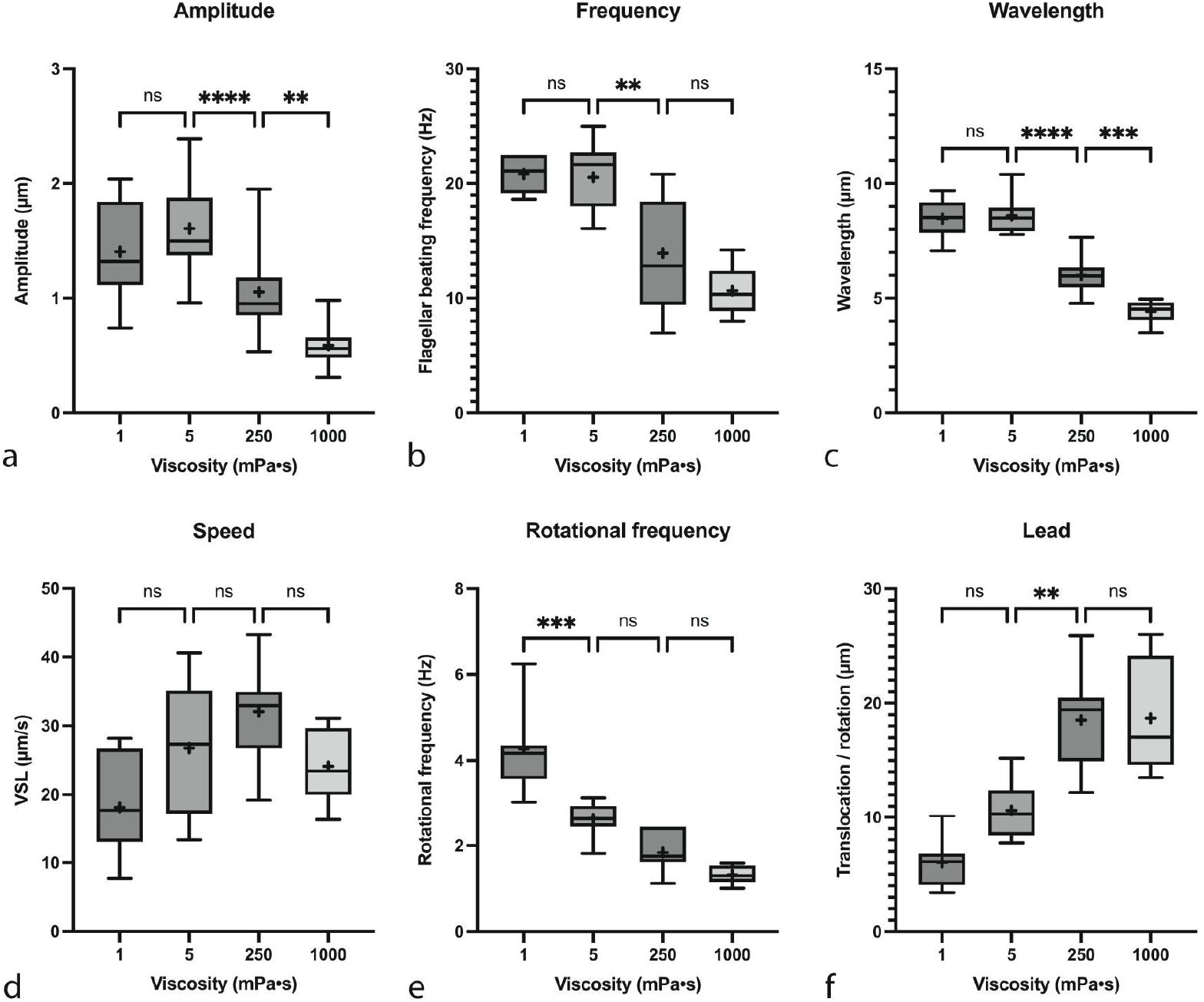
Quantification of *T. brucei* MiTat 1.6 forward swimming parameters. **a)** Amplitude of the anterior-most wavelength as measured during beating in the image plane (white lines in Fig. 1). **b)** Flagellar beating frequency measured as number of anterior-initiated waves during time periods of uninterrupted swimming. **c)** Wavelength of the anterior flagellum (blue lines in Fig.1). **d)** Swimming speed, measured as straight line velocity of the posterior tip (dashed white lines in Fig.1). **e)** Rotational frequency, measured as the number of uninterrupted 360° rotations around the longitudinal axis per time unit. **f)** Lead, measured as the translocation per 360° rotation (dashed white lines in Fig. 1). Box plots show the mean (+) and median (line) values within the 25th to 75th percentile range as boxes and the min to max values as whiskers (n=7 cells and 2-4 rotations for each parameter). Significance was analysed by ordinary one-way ANOVA (****: p < 0,0001, ***: p<0.001, **: p<0.01, *: p<0.05, ns: p>0.05).

**Video S1** One rotation of a persistently swimming trypanosome at 1 mPa*s. Cell analysed in Fig. 1b, 120 ms, 25x slowed.

**Video S2** One rotation of a persistently swimming trypanosome at 5 mPa*s Cell analysed in Fig. 1c, 220 ms, 25x slowed.

**Video S3** One rotation of a persistently swimming trypanosome at 250 mPa*s Cell analysed in Fig. 1d, 760 ms, 25x slowed.

**Video S4** One rotation of a persistently swimming trypanosome at 1000 mPa*s Cell analysed in Fig. 1e, 980 ms, 25x slowed.

**Video S5** Direct comparison of several rotations of persistently swimming trypanosomes in different viscosities Cells used for trajectories in Fig. 3, 10x slowed.

**Video S6** MiTat 1.6 cell swimming in 1mPa*s through pillars with 8µm diameter and 4µm spacing.

**Linked html file 1** Rotatable 3D-plot corresponding to Fig. 4a2

**Linked html file 2** Rotatable 3D-plot corresponding to Fig. 4b2

**Linked html file 3** Rotatable 3D-plot corresponding to Fig. 5a

**Linked html file 4** Rotatable 3D-plot corresponding to Fig. 5b

**Video S7** Trypanosome swimming in 1000 mPa*s viscosity. The cell body does not rotate, therefore the swimming trajectory is circular. Cell shown in Fig. 6a, 4x slowed.

**Video S8** Trypanosome switching from rotational to non-rotational circular swimming in 1000 mPa*s (interrupted by several backward beats and short reverse movement). Cell shown in Fig. 6b, 4x slowed.

**Video S9** Trypanosome motile behaviour in 4000 mPa*s viscosity. Cell shown in Fig. 7a, original speed.

**Video S10** Trypanosome motile behaviour in collagen gel (4%). Cell shown in Fig. 7b, original speed.

## References

1. Cayla M, Rojas F, Silvester E, Venter F, Matthews KR. African trypanosomes. Parasit Vectors. 2019 Dec;12(1):190.

2. Overberg FA, Khameneh NJ, Krüger T, Engstler M, Gompper G, Fedosov DA. Modelling motility of Trypanosoma brucei. PLOS Comput Biol. 2025 May 21;21(5):e1013111.

3. Tan Z, Peters JIU, Stark H. Trypanosoma brucei moving in microchannels and through constrictions. New J Phys. 2025 May;27(6):064401.

4. Krüger T, Schuster S, Engstler M. Beyond Blood: African Trypanosomes on the Move. Trends Parasitol. 2018 Dec;34(12):1056–67.

5. Krüger T, Engstler M. Trypanosomes – versatile microswimmers. Eur Phys J Spec Top. 2016 Nov 1;225(11–12):2157–72.

6. Heddergott N, Krüger T, Babu SB, Wei A, Stellamanns E, Uppaluri S, et al. Trypanosome Motion Represents an Adaptation to the Crowded Environment of the Vertebrate Bloodstream. PLoS Pathog. 2012 Nov;8(11):e1003023.

7. Elgeti J, Winkler RG, Gompper G. Physics of microswimmers—single particle motion and collective behavior: a review. Rep Prog Phys. 2015 May 1;78(5):056601.

8. Crilly NP, Mugnier MR. Thinking outside the blood: Perspectives on tissue-resident Trypanosoma brucei. PLoS Pathog. 2021 Sept 16;17(9):e1009866.

9. Schuster S, Krüger T, Subota I, Thusek S, Rotureau B, Beilhack A, et al. Developmental adaptations of trypanosome motility to the tsetse fly host environments unravel a multifaceted in vivo microswimmer system. eLife. 2017 Aug 15;6:e27656.

10. Bargul JL, Jung J, McOdimba FA, Omogo CO, Adung’a VO, Krüger T, et al. Species-Specific Adaptations of Trypanosome Morphology and Motility to the Mammalian Host. Carrington M, editor. PLOS Pathog. 2016 Feb 12;12(2):e1005448.

11. Beaver AK, Keneskhanova Z, Cosentino RO, Weiss BL, Awuoche EO, Smallenberger GM, et al. Tissue spaces are reservoirs of antigenic diversity for Trypanosoma brucei. Nature. 2024 Dec;636(8042):430–7.

12. Carvalho T, Trindade S, Pimenta S, Santos AB, Rijo-Ferreira F, Figueiredo LM. Trypanosoma brucei triggers a marked immune response in male reproductive organs. PLoS Negl Trop Dis. 2018 Aug 15;12(8):e0006690.

13. Coles JA, Myburgh E, Ritchie R, Hamilton A, Rodgers J, Mottram JC, et al. Intravital Imaging of a Massive Lymphocyte Response in the Cortical Dura of Mice after Peripheral Infection by Trypanosomes. PLoS Negl Trop Dis. 2015 Apr 16;9(4):e0003714.

14. Mabille D, Dirkx L, Thys S, Vermeersch M, Montenye D, Govaerts M, et al. Impact of pulmonary African trypanosomes on the immunology and function of the lung. Nat Commun. 2022 Nov 18;13(1):7083.

15. Mogk S, Meiwes A, Shtopel S, Schraermeyer U, Lazarus M, Kubata B, et al. Cyclical Appearance of African Trypanosomes in the Cerebrospinal Fluid: New Insights in How Trypanosomes Enter the CNS. PLoS ONE. 2014 Mar 11;9(3):e91372.

16. Reuter C, Hauf L, Imdahl F, Sen R, Vafadarnejad E, Fey P, et al. Vector-borne Trypanosoma brucei parasites develop in artificial human skin and persist as skin tissue forms. Nat Commun. 2023 Nov 23;14(1):7660.

17. Trindade S, Rijo-Ferreira F, Carvalho T, Pinto-Neves D, Guegan F, Aresta-Branco F, et al. Trypanosoma brucei Parasites Occupy and Functionally Adapt to the Adipose Tissue in Mice. Cell Host Microbe. 2016 June 8;19(6):837–48.

18. Trindade S, De Niz M, Costa-Sequeira M, Bizarra-Rebelo T, Bento F, Dejung M, et al. Slow growing behavior in African trypanosomes during adipose tissue colonization. Nat Commun. 2022 Dec 8;13(1):7548.

19. Wolburg H, Mogk S, Acker S, Frey C, Meinert M, Schönfeld C, et al. Late Stage Infection in Sleeping Sickness. PLOS ONE. 2012 Mar 27;7(3):e34304.

20. Machado H, Temudo A, Niz MD. The lymphatic system favours survival of a unique T. brucei population. Biol Open. 2023 Nov 9;12(11):bio059992.

21. Capewell P, Cren-Travaillé C, Marchesi F, Johnston P, Clucas C, Benson RA, et al. The skin is a significant but overlooked anatomical reservoir for vector-borne African trypanosomes. eLife. 2016;5:e17716.

22. Swartz MA, Fleury ME. Interstitial Flow and Its Effects in Soft Tissues. Annu Rev Biomed Eng. 2007 Aug 15;9(Volume 9, 2007):229–56.

23. Lauga E. Bacterial Hydrodynamics. Annu Rev Fluid Mech. 2016 Jan 3;48(Volume 48, 2016):105–30.

24. Lauga E, Powers TR. The hydrodynamics of swimming microorganisms. Rep Prog Phys. 2009 Sept 1;72(9):096601.

25. Kamdar S, Shin S, Leishangthem P, Francis LF, Xu X, Cheng X. The colloidal nature of complex fluids enhances bacterial motility. Nature. 2022 Mar 31;603(7903):819–23.

26. Li G, Lauga E, Ardekani AM. Microswimming in viscoelastic fluids. J Non-Newton Fluid Mech. 2021 Nov 1;297:104655.

27. Spagnolie SE, Underhill PT. Swimming in Complex Fluids. Annu Rev Condens Matter Phys. 2023;14(1):381–415.

28. Büyükurgancı B, Basu SK, Neuner M, Guck J, Wierschem A, Reichel F. Shear rheology of methyl cellulose based solutions for cell mechanical measurements at high shear rates. Soft Matter. 2023 Mar 1;19(9):1739–48.

29. Ishimoto K, Gadêlha H, Gaffney EA, Smith DJ, Kirkman-Brown J. Human sperm swimming in a high viscosity mucus analogue. J Theor Biol. 2018 June;446:1–10.

30. Vassella E, Boshart M. High molecular mass agarose matrix supports growth of bloodstream forms of pleomorphic Trypanosoma brucei strains in axenic culture. Mol Biochem Parasitol. 1996 Nov 12;82(1):91–105.

31. Krüger T, Engstler M. Motility Analysis of Trypanosomatids. In: Michels PAM, Ginger ML, Zilberstein D, editors. Trypanosomatids [Internet]. New York, NY: Springer US; 2020 [cited 2020 June 10]. p. 409–23. (Methods in Molecular Biology; vol. 2116). Available from: http://link.springer.com/10.1007/978-1-0716-0294-2_25

32. Gray J, Hancock GJ. The Propulsion of Sea-Urchin Spermatozoa. J Exp Biol. 1955 Dec 1;32(4):802–14.

33. Hancock GJ. The self-propulsion of microscopic organisms through liquids. Proc R Soc Lond Ser Math Phys Sci. 1953;217(1128):96–121.

34. Lighthill J. Flagellar Hydrodynamics. SIAM Rev. 1976 Apr;18(2):161–230.

35. Engstler M, Pfohl T, Herminghaus S, Boshart M, Wiegertjes G, Heddergott N, et al. Hydrodynamic Flow-Mediated Protein Sorting on the Cell Surface of Trypanosomes. Cell. 2007 Nov 2;131(3):505–15.

36. Wheeler RJ. Use of chiral cell shape to ensure highly directional swimming in trypanosomes. Rao CV, editor. PLOS Comput Biol. 2017 Jan 31;13(1):e1005353.

37. Wróbel JK, Lynch S, Barrett A, Fauci L, Cortez R. Enhanced flagellar swimming through a compliant viscoelastic network in Stokes flow. J Fluid Mech. 2016 Apr;792:775–97.

38. Gong A, Rode S, Gompper G, Kaupp UB, Elgeti J, Friedrich BM, et al. Reconstruction of the three-dimensional beat pattern underlying swimming behaviors of sperm. Eur Phys J E. 2021 July 1;44(7):87.

39. Ren X, Bloomfield-Gadêlha H. Swimming by Spinning: Spinning-Top Type Rotations Regularize Sperm Swimming Into Persistently Progressive Paths in 3D. Adv Sci. 2025;12(6):2406143.

40. Gadadhar S, Alvarez Viar G, Hansen JN, Gong A, Kostarev A, Ialy-Radio C, et al. Tubulin glycylation controls axonemal dynein activity, flagellar beat, and male fertility. Science. 2021 Jan 8;371(6525):eabd4914.

41. Friedrich BM, Riedel-Kruse IH, Howard J, Julicher F. High-precision tracking of sperm swimming fine structure provides strong test of resistive force theory. J Exp Biol. 2010 Apr 15;213(8):1226–34.

42. Elgeti J, Kaupp UB, Gompper G. Hydrodynamics of Sperm Cells near Surfaces. Biophys J. 2010 Aug 9;99(4):1018–26.

43. Chaudhuri O, Cooper-White J, Janmey PA, Mooney DJ, Shenoy VB. Effects of extracellular matrix viscoelasticity on cellular behaviour. Nature. 2020 Aug;584(7822):535–46.

44. Cuche E, Bevilacqua F, Depeursinge C. Digital holography for quantitative phase-contrast imaging. Opt Lett. 1999 Mar 1;24(5):291–3.

